# Brain neurosteroids are natural anxiolytics targeting α2 subunit γ-aminobutyric acid type-A receptors

**DOI:** 10.1101/462457

**Authors:** Elizabeth J Durkin, Laurenz Muessig, Tanja Herlt, Michael J Lumb, Ryan Patel, Philip Thomas, Damian P. Bright, Rachel Jurd, Stephen J Moss, Anthony H. Dickenson, Francesca Cacucci, Trevor G Smart

## Abstract

Neurosteroids are naturally-occurring molecules in the brain that modulate neurotransmission. They are physiologically important since disrupting their biosynthesis precipitates neurological disorders, such as anxiety and depression. The endogenous neurosteroids, allopregnanolone and tetrahydro-deoxycorticosterone are derived from sex and stress hormones respectively, and exhibit therapeutically-useful anxiolytic, analgesic, sedative, anticonvulsant and antidepressant properties. Their main target is the γ-aminobutyric acid type-A inhibitory neurotransmitter receptor (GABA_A_R), whose activation they potentiate. However, whether specific GABA_A_R isoforms and neural circuits differentially mediate endogenous neurosteroid effects is unknown. By creating a knock-in mouse that removes neurosteroid potentiation from α2-GABA_A_R subunits, we reveal that this isoform is a key target for neurosteroid modulation of phasic and tonic inhibition, and is essential for the anxiolytic role of endogenous neurosteroids, but not for their anti-depressant or analgesic properties. Overall, α2-GABA_A_R targeting neurosteroids may act as selective anxiolytics for the treatment of anxiety disorders, providing new therapeutic opportunities for drug development.

## Introduction

GABA_A_Rs are crucial moderators of neuronal excitability in the CNS where the relative efficacy of GABA-mediated inhibition will depend, at least in part, on the positive allosteric modulation of such receptors by endogenous neurosteroids. The levels of two major neurosteroids in the brain, allopregnanolone and THDOC, are increased during pregnancy (*Luisi et al., 2000*; *Concas et al., 1998*) and in response to stressors (*Purdy et al., 1991*; *Paul and Purdy, 1992*; *Barbaccia et al., 1998*) or drugs such as nicotine (*Porcu et al., 2003*), γ-hydroxybutyrate (*Barbaccia et al., 2002*) and some antidepressants (*Uzunov et al., 1996*). Increased levels of endogenous neurosteroids may also underlie the antidepressant and anticonvulsant effects of ethanol (*Helms et al., 2012*; *Hirani et al., 2002*; *Khisti et al., 2002*; *VanDoren et al., 2000*). By comparison, dysfunctional neurosteroid production is frequently associated with a range of diseases, including premenstrual dysphoric disorder, panic disorder, anxiety and stress, depression, schizophrenia, bipolar disorder, eating disorders, and dementia (*Mellon and Griffin, 2002*; *Strous et al., 2006*).

To explore the therapeutic potential of neurosteroids, it is necessary to first define their range of physiological actions at molecular, cellular and systems levels. Their most important target is the GABA_A_R (*Belelli et al., 2006*), at which they potentiate receptor activation by binding to a site located at the receptor’s β-α subunit interface (*Hosie et al., 2006*; *Laverty et al., 2017*; *Miller et al., 2017*). This potentiation site is highly conserved across the GABA_A_R family and a key residue in the α subunit plays a critically important role in binding (*Hosie et al., 2009*). Thus GABA_A_Rs containing many combinations of αβγ or αβδ subunits could, in principle, mediate the physiological effects of neurosteroids. We hypothesized that the actions of neurosteroids may segregate with different α subunit isoforms (α1-α6), as noted previously for the benzodiazepines, for which α1-containing GABA_A_Rs mediate sedation, amnesia and anticonvulsion (*Rudolph et al., 1999*; *McKernan et al., 2000*), whilst α2 and α3-containing GABA_A_Rs confer their anxiolytic and myorelaxant effects (*Low et al., 2000*). If α2-type GABA_A_Rs are involved in the anxiolytic effects of endogenous neurosteroids, then α2-subunit-selective neurosteroid analogues should behave as non-sedating anxiolytics, potentially lacking deleterious drug-withdrawal effects (*Rupprecht et al., 2009*). This outcome could facilitate new drug development opportunities with advantages over the widely-used benzodiazepines.

To examine the physiological roles of neurosteroids, we created a knock-in mouse line by replacing the hydrogen-bonding glutamine at position 241 located in the first transmembrane domain of α2 subunits, which is pivotal to neurosteroid sensitivity (*Hosie et al., 2006*). Substitution with hydrophobic methionine (α2^Q241M^) ablated this sensitivity, enabling the physiological roles played by endogenous positive allosteric neurosteroids when modulating α2-GABA_A_Rs to be deduced and explored with precision. The choice of this subunit was also influenced by the physiological importance of α2-containing GABA_A_Rs in controlling neuronal spike output profiles (*Nusser et al., 1996*), due to their clustered localization at the axon initial segment, and their abundant expression in the hippocampus and medial prefrontal cortex (mPFC) (*Laurie et al., 1992b*; *Laurie et al., 1992a*; *Wisden et al., 1992*; *Pirker et al., 2000*), where they may contribute significantly to synaptic inhibition (*Prenosil et al., 2006*).

Characterisation of this knock-in mouse line revealed that synaptic α2-GABA_A_Rs are clearly sensitive to modulation by endogenous neurosteroids. We also discovered that neurosteroid potentiation of tonic currents in the dentate gyrus was absent in the knock-in mice, highlighting an unexpected contribution of α2 subunits to extrasynaptic GABA-mediated inhibition. At a systems level, previous work has demonstrated that knock-out of the α2 subunit caused mice to become more anxious in a conditioned emotional response paradigm (*Dixon et al., 2008*). Here, we demonstrate an anxiety-like phenotype in our α2 subunit knock-in line, providing unequivocal evidence for endogenous neurosteroids having a basal anxiolytic effect in an unperturbed animal, mediated via α2-type GABA_A_Rs and their associated neural circuits.

## Results

### α2^Q241M^ knock-in does not affect GABA_A_ receptor subunit expression

The point mutation α2^Q241M^ specifically ablates neurosteroid modulation at recombinant α2β2/3γ2 GABA_A_Rs without affecting GABA sensitivity, receptor activation or the actions of other allosteric modulators at these receptors (*Hosie et al., 2006*; *Hosie et al., 2009*). We therefore generated a mouse line with this mutation, using a targeting vector containing the point mutation α2^Q241M^ in exon 8 (*figure supplement 1*), which was electroporated into embryonic stem cells prior to blastocyst injection and breeding. The presence of mutant alleles was subsequently confirmed by Southern blotting and sequence analyses (*Figure 1a,b*).

**Figure 1.**
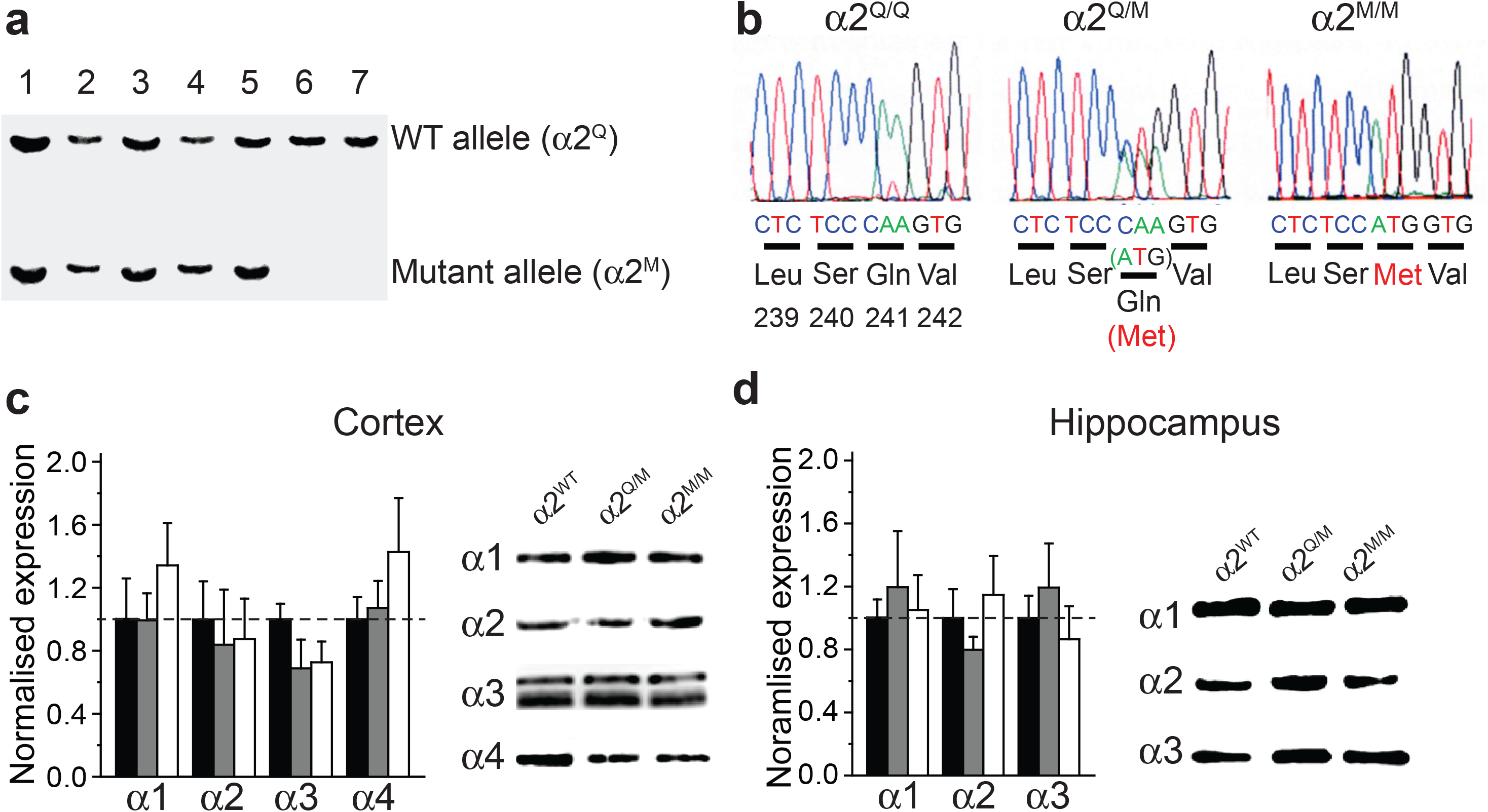
Creating and validating a neurosteroid binding site α2^Q241M^ knock-in mouse line. **(a)** Southern blot screen of ES cell lines: the Q241M mutation silently introduces a novel restriction site – allowing a screen for restriction fragment length polymorphism. Lanes 1 - 5, heterozygous recombinants; lanes 6 - 7, wild-type littermate controls. **(b)** Verifying the point mutation by DNA sequencing. Exon 8 was PCR-amplified from mouse genomic DNA of wild-type (α2^Q/Q^: left), α2^Q/M^ (middle) and α2^M/M^ (right) animals, then sequenced. Sequence data around the point mutation are shown. **(c,d)** GABA_A_R α1-α4 subunit expression levels were determined by Western blotting total protein isolated from the cortex (**c**) and hippocampus (**d**). Left panels, quantitation of expression in α2^Q/M^ (grey) and α2^M/M^ (white) relative to wild-type (α2^Q/Q^, black) brain samples. Right panels, representative Western blots from cortex and hippocampus for wild-type (WT), α2^Q/M^ and α2^M/M^ mice. Images show blots between 45 and 66 kDa markers. Loading controls with tubulin are omitted for clarity. Data are mean ± sem, one-way ANOVA, n = 6-9. There are no statistically significant differences between groups.

The knock-in mutation did not cause any adverse phenotypes, and homozygote (α2^M/M^) and heterozygote (α2^Q/M^) knock-ins, and wild-type (α2^Q/Q^) littermates, were produced in approximately Mendelian ratios. Using a combination of quantitative Western blotting (*Figure 1c,d*) and immunohistochemistry (*figure supplements 2-4*) we assessed the expression patterns of GABA_A_R α1 - α5 subunits in the cortex and hippocampus. This revealed no evidence of any compensatory effects as a consequence of the α2 knock-in mutation. Thus, the loss of basal neurosteroid potentiation at α2-GABA_A_Rs does not re-configure the expression pattern for GABA_A_R α subunits. Consequently, any phenotypes of the knock-in mice can be attributed directly to the loss of neurosteroid potentiation at α2-GABA_A_Rs.

### GABA-mediated synaptic inhibition in α2^Q241M^ hippocampal and prefrontal cortical slices

The functional effects of ablating the α2 subunit neurosteroid binding site were assessed on synaptic inhibition using whole-cell recording from hippocampal CA1 pyramidal neurons (PNs) and dentate gyrus granule cells (DGGCs), and from layer II/III pyramidal neurons in the medial prefrontal cortex (mPFC) in acute brain slices. These brain regions strongly express α2-GABA_A_Rs (*Pirker et al., 2000*; *Sperk et al., 1997*) that are typically associated with inhibitory synapses and rapid phasic inhibition (*Prenosil et al., 2006*) and moreover, they contain inhibitory circuitry thought to be important for the regulation of anxiety-motivated behaviours (*Engin and Treit, 2007*; *Engin et al., 2016*).

Initially, we examined whether synaptic α2-GABA_A_Rs were responsive to basal endogenous levels of neurosteroids that should be revealed by genotypic differences in baseline GABAergic neurotransmission and the kinetics of spontaneous inhibitory postsynaptic currents (sIPSCs). Typically, spontaneous IPSCs (sIPSCs) respond to neurosteroids with prolonged weighted decay times (τ_w_) (*Otis and Mody, 1992*; *Nusser and Mody, 2002*; *Belelli and Herd, 2003*; *Harney et al., 2003*). Interestingly, sIPSCs from α2^M/M^ mice decayed at significantly faster rates (by up to ~20 – 30 %) compared to synaptic events recorded from wild-type littermates (CA1 PNs and DGGCs (*Figure 2a*); mPFC PNs (*figure supplement 5*), whilst sIPSC rise-times, amplitudes and frequencies remained unaltered (*table supplement 1*).

**Figure 2.**
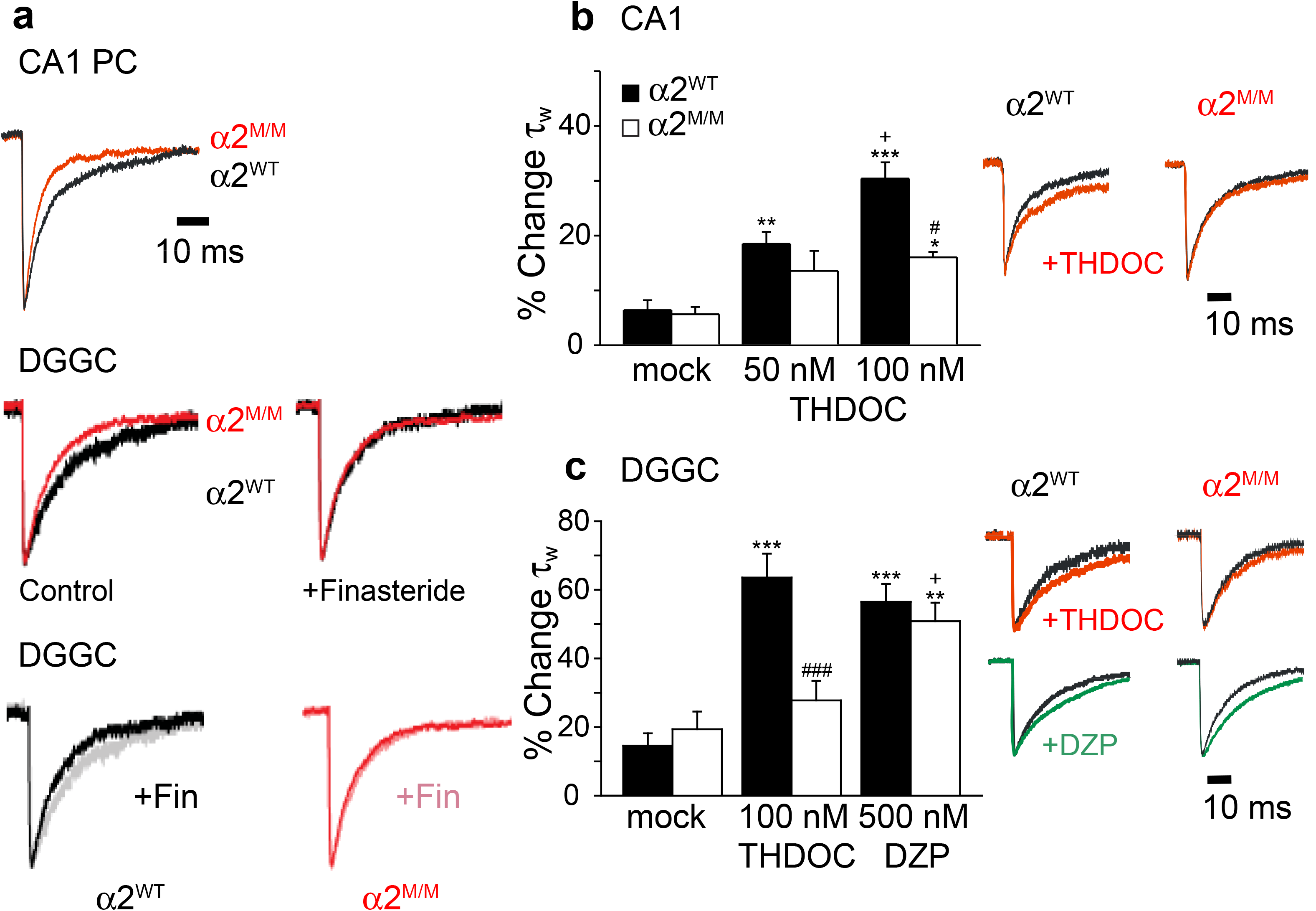
Synaptic currents in α2^Q241M^ knock-in hippocampal slices. **(a)** Peak-scaled averaged sIPSCs recorded from: Top, CA1 principal cells (PC) for wild-type (black) and α2^M/M^ (red) hippocampal slices; Middle, DGGCs, to reveal the effect of genotype (wild-type (black), α2^M/M^ (red)) on sIPSC decay times under control conditions (left panel), and after treatment with 50 μM finasteride (right panel); Bottom, DGGCs, demonstrating how control sIPSC decays for wild type (grey) and α2^M/M^ (red) are differentially affected by finasteride pre-treatment (black and pink respectively). (**b and c)** Bar charts for THDOC and diazepam (DZP) effects on weighted decay times (τ_w_) for sIPSCs recorded from CA1 neurons (**b**) and DGGCs (**c**) for wild-type (black) and α2^M/M^ (white) slices. Insets show representative peak-scaled averaged sIPSCs for wild-type and α2^M/M^ cells for control epochs (black) and after addition of 100 nM THDOC (red) or 500 nM diazepam (green). Data are mean ± sem. Statistical significance indicated as: * p<0.05, ** p<0.01, and *** p<0.001 for the effects of THDOC or diazepam compared to mock/vehicle; + p<0.05 for diazepam vs. THDOC, and for effect of 100 nM THDOC vs. 50 nM THDOC; # p<0.05 and ### p<0.001 for effect of genotype (i.e. α2^M/M^ vs. wild-type). See Table supplements 1-3 for statistical analyses.

We then examined sIPSC kinetics in DGGCs in the absence of endogenous neurosteroids, by pre-incubating hippocampal slices with the neurosteroid synthesis inhibitor, finasteride, which blocks the 5α-reductase enzyme and can affect the profiles of sIPSCs in brain slices (*Keller et al., 2004*). Interestingly, the genotypic difference evident in the weighted decay times for sIPSCs recorded from DGGCs for wild-type and α2^M/M^ littermates was abolished by finasteride (50 μM; *Figure 2a - table supplement 1*). Thus, removing endogenous neurosteroids significantly reduced τ_w_ in DGGCs from wild-type animals, but had no effect on τ_w_ in α2^M/M^ mice. These data revealed the presence of an endogenous neurosteroid tone within the hippocampus, and highlighted the prominent role played by synaptic α2-GABA_A_Rs in responding to neurosteroids, thereby setting an increased level of basal inhibition in this brain region. The lack of effect of finasteride in α2^M/M^ mice further emphasised not only the absence of compensation for the loss of the α2 subunit neurosteroid binding site, but also the importance of α2-GABA_A_Rs in DGGCs in mediating the basal effects of neurosteroids (*Figure 2a*).

We next assessed the response of synaptic GABA currents to an applied neurosteroid (THDOC). In slices from WT mice, THDOC increased the duration of sIPSCs in CA1 PNs, DGGCs and mPFC PNs by increasing τ_w_ without changing sIPSC amplitudes, rise-times or frequencies (*Figure 2b,c – figure supplement 5 – table supplements 2-4*). In contrast, and as predicted, sIPSC prolongation by 100 nM THDOC was significantly diminished in cells from α2^M/M^ mice compared to those from wild-type littermates (*Figure 2b,c; figure supplement 5; table supplements 2-4*) consistent with a reduced potency and/or efficacy of THDOC. Given that the Q241M mutation only prevents neurosteroid potentiation at α2-GABA_A_Rs, the residual neurosteroid modulation of synaptic GABA currents observed in both CA1 and mPFC PNs of α2^M/M^ slices (*Figure 2b; figure supplement 5*) can only be attributed to other α subunit receptor isoforms that are expressed in these neurons (e.g. α1, α3, α4 and α5) (*Pirker et al., 2000*). For DGGCs there was less evidence of residual modulation (*Figure 2c*). We conclude that on the basis of differential THDOC modulation of sIPSCs in wild-type and α2^M/M^ neurons (*Figure 2b,c – figure supplement 5*), α2-GABA_A_Rs contribute significantly to neurosteroid regulation of synaptic inhibition in CA1 and layer II/II mPFC PNs and especially DGGCs.

The α2^Q241M^ mutation was predicted to specifically reduce modulation of sIPSC by neurosteroids, but not by benzodiazepines, which bind at a distinct site on synaptic αβγ-containing GABA_A_Rs (*Hosie et al., 2006*; *Sigel and Steinmann, 2012*). To test this prediction, we assessed the effect of diazepam on sIPSCs in DGGCs. In contrast to the reduced THDOC effect, the diazepam-induced prolongation of τ_w_ was unaffected by the α2^Q241M^ mutation compared to wild-type (*Figure 2c – table supplement 3*). Therefore, the α2^Q241M^ mutation specifically ablates the response to neurosteroids, and does not initiate compensatory changes that diminish synaptic GABA_A_R sensitivity to other positive allosteric modulators.

### α2 subunit GABA_A_ receptors contribute to tonic inhibition in the hippocampus

We used the neurosteroid binding site mutation to assess if α2-GABA_A_Rs residing in the extrasynaptic domain can contribute towards GABA-mediated tonic inhibition. The consensus view of extrasynaptic GABA_A_Rs underlying tonic inhibition in the hippocampus and cortex is that they are most likely composed of α4βδ, α5βγ and α1β2, with no expectation of a contribution from α2-GABA_A_Rs (*Farrant and Nusser, 2005*). We focussed on DGGCs and mPFC PNs as only very small tonic currents were evident in CA1 PNs. In the cortical neurons, there was no difference in basal tonic currents recorded from wild-type and α2^M/M^ slices, and application of 100 nM THDOC caused comparable potentiation of the tonic current in both genotypes (*table supplement 6*). These results indicate that neurosteroid-sensitive α2-GABA_A_Rs are unlikely to contribute to tonic inhibition in layer II/II mPFC PNs.

Similarly, there was no effect of α2^Q241M^ on the basal tonic current in DGGCs (*table supplement 5*). Application of 100 nM THDOC caused a significant inward shift in the holding current for wild-type DGGCs (*Figure 3a,b*), which consequently led to a large increase in the bicuculline-sensitive tonic current compared to control (139 % increase, *table supplement 5*). Unexpectedly, the increased holding current induced by neurosteroid was significantly reduced by ~5-fold in DGGCs from α2^M/M^ slices (27% increase, *table supplement 5*), suggesting that the THDOC potentiation of tonic inhibition must involve a significant α2-containing GABA_A_R population. Furthermore, 500 nM diazepam had no modulatory effect on the bicuculline-sensitive tonic current (*Figure 3a*), indicating that the GABA_A_Rs mediating the tonic current are mostly devoid of the γ subunit. These observations represent the first direct evidence of a new role for α2 subunit-containing GABA_A_Rs, typically thought to be associated only with receptors at inhibitory synapses

**Figure 3.**
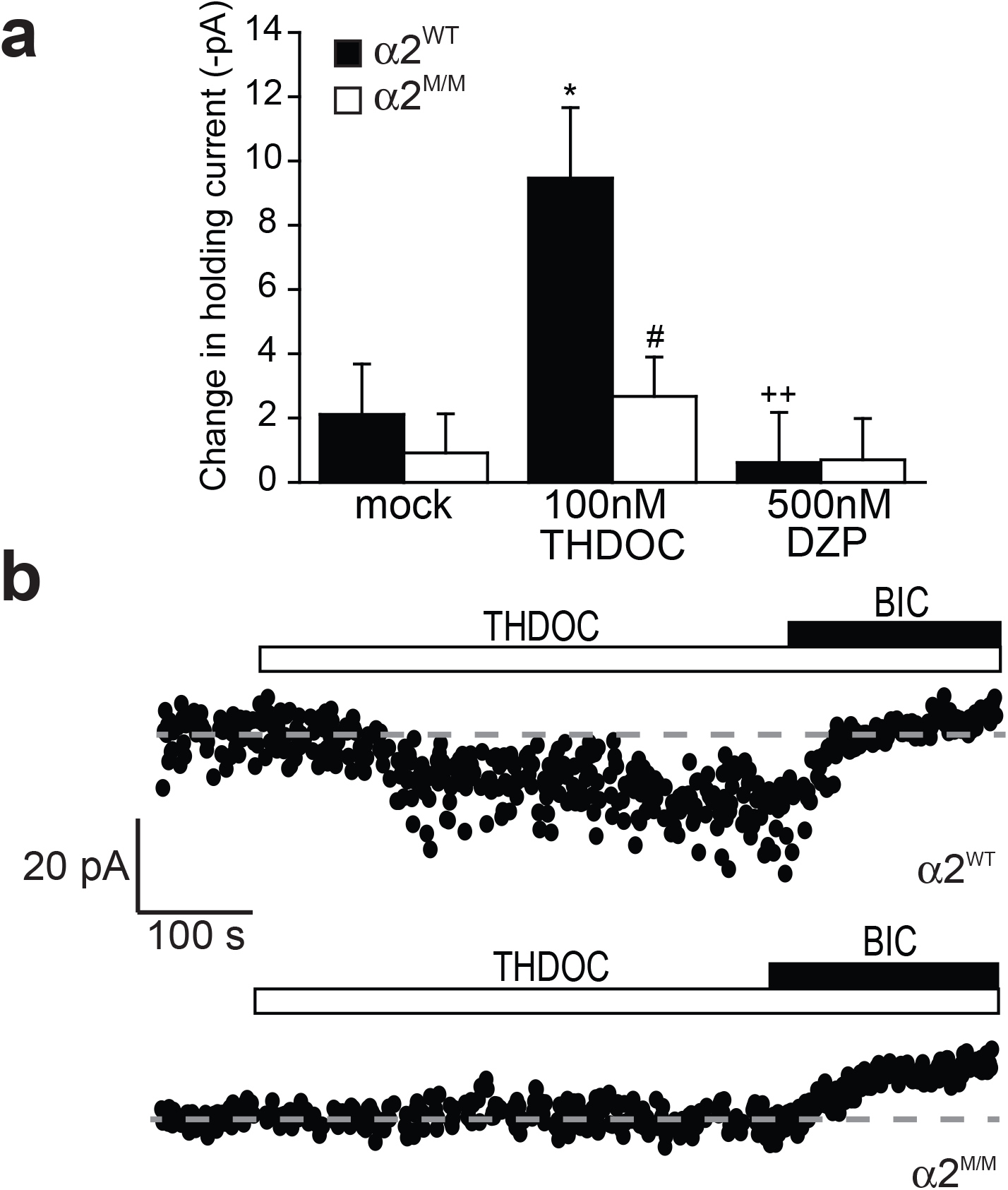
Tonic inhibition in α2^Q241M^ knock-in hippocampal slices. **(a)** Bar chart representing changes in the holding current (-pA) for DGGCs from wild-type (black) and α2^M/M^ (white) after application of DMSO (mock), THDOC (100 nM) or diazepam (DZP, 500 nM). (**b)** Holding currents from wildtype and α2^M/M^ DGGCs under control conditions, and during the application of: 100 nM THDOC (open bar); and 20 μM bicuculline (black bar). Data in (a) are mean ± sem. Statistical significance indicated as: *p <0.05 for the effects of THDOC compared to mock/DMSO; ^++^ p<0.01 for the effect of diazepam vs THDOC; ^#^ p<0.05 for the effect of genotype (α2^M/M^ vs wild-type).

(*Tretter et al., 2008*), in mediating extrasynaptic tonic GABA currents. We speculate that these receptors are likely to be composed of α2β or α2βδ subunits. Such unusual extrasynaptic receptors are readily assembled by co-expression of α2, β2 and δ subunits in HEK293 cells. Furthermore, the Q241M mutation prevented the neurosteroid modulation that is manifest by a small increase in GABA potency and a larger increase in GABA macroscopic efficacy by ~50 % for the α2βδ receptors (*figure supplement 6 – table supplement 7*). This effect of neurosteroids contrasts with their modulation of α2β2γ2L or α2β3γ2L receptors where only increases in GABA potency were noted, which are ablated by Q241M (*figure supplement 6 – table supplement 7*).

### Anxiogenic phenotype caused by removing neurosteroid sensitivity from α2-GABA_A_Rs

The physiological importance and role of neurosteroid modulation of α2-GABA_A_Rs was assessed *in vivo* using an array of behavioural tests. Anxiety-based phenotypes were explored using three benchmark screens: the elevated plus maze (EPM), the light-dark box (LDB), and suppression of feeding behaviour for novel food (hyponeophagia). If the physiological role(s) of neurosteroids mediated via α2-GABA_A_Rs involve anxiolysis, then α2^M/M^ mice might be expected to show a baseline phenotype implicating α2-GABA_A_Rs and basal neurosteroid tone in modulating anxiety-based behaviours. Furthermore, the response to injected neurosteroids should also be affected in the α2^M/M^ mice.

We used the EPM to establish basal anxiety-like phenotypes, and observed that α2^M/M^ mice spent significantly less time on and tended to make fewer entries into the open arms compared to wild-type littermates (*Figure 4a,b*). This behaviour was not caused by differences in locomotor activity since entries into closed arms, as well as total arm entries (*Figure 4c – figure supplement 7*), which are indicators of activity, remained unaltered on comparing α2^WT^ with α2^M/M^. The anxiety phenotype for α2^M/M^ mice over wild-type littermates was also evident using the LDB, in which α2^M/M^ mice were not only quicker to leave, but also spent less time in the aversive light zone and thus more time in the dark zone (*Figure 4d,e*). In addition, the α2^M/M^ mice made fewer transfers between the light and dark zones (*Figure 4f*), and showed less exploratory activity in the dark zone (*Figure 4g*) compared to wild-type. The heightened basal anxiety evident in the α2^M/M^ line was further corroborated using the hyponeophagia test. Rodents will initially suppress feeding behaviour when encountering novel food, even if the food is appealing for consumption and the animal is also hungry. The latency to approach and consume novel foods is an indicative marker of anxiety. For the α2^M/M^ mice this latency was markedly increased compared to that for α2^WT^ (*Figure 4h*). Taken together, all these observations are consistent with a more anxious phenotype for the a2^M/M^ mouse and suggest that endogenous neurosteroids perform a physiological role managing or reducing baseline anxiety by potentiating inhibition mediated by α2-GABA_A_Rs.

**Figure 4.**
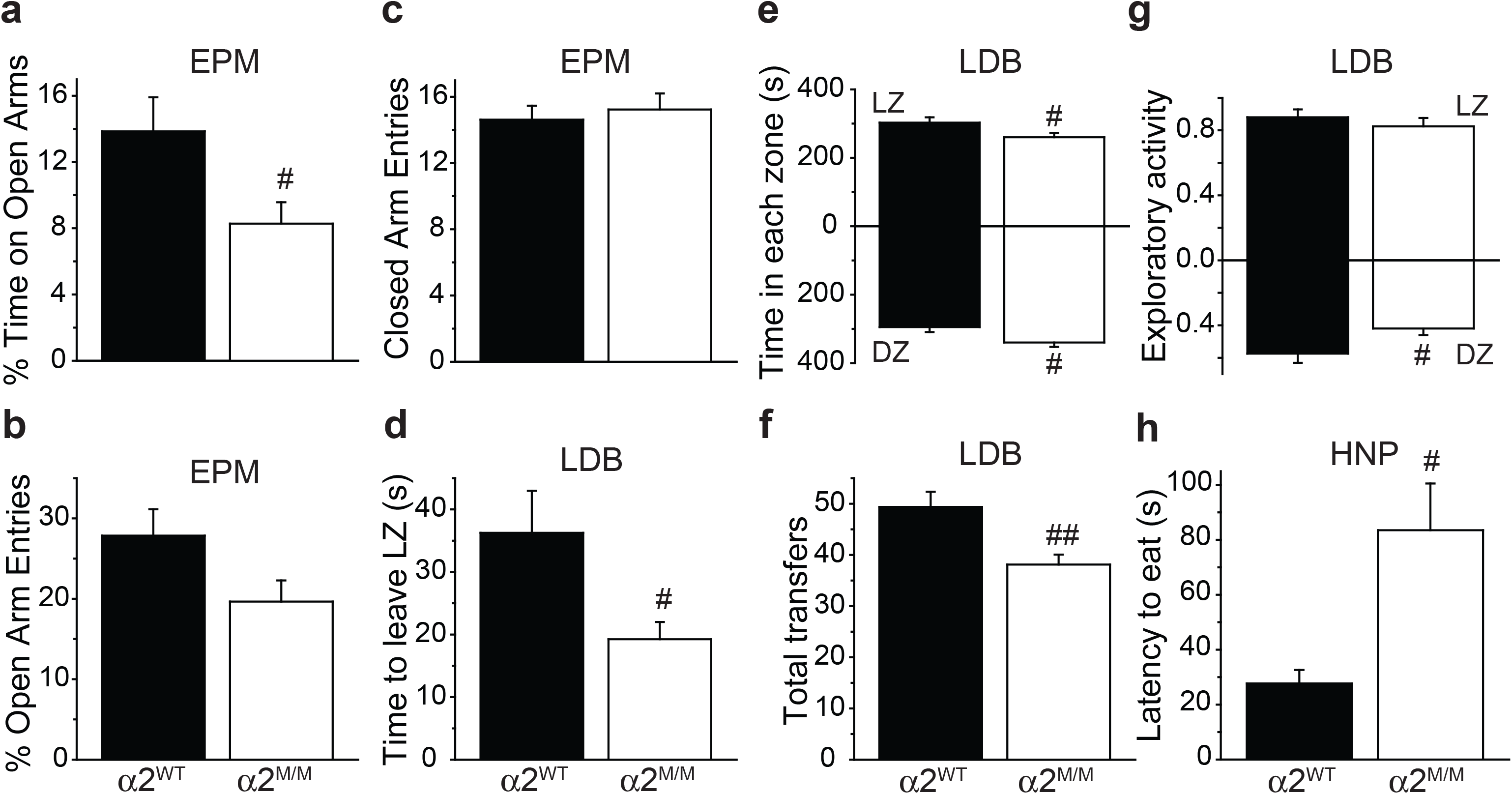
Anxiogenic phenotype of α2^Q241M^ knock-in. **(a and b)** α2^M/M^ mice (white) spent less time on the open arms of the elevated plus maze (EPM) than α2^WT^ (black) mice and tend to make fewer entries into the open arms. This difference is not due to differences in locomotor activity as closed arm entries (**c**) are unchanged. (**d-g)** In the light-dark box (LDB), α2^M/M^ mice (white) were faster to leave the light zone (LZ) (**d**), spent less time in the light zone and more time in the dark zone (**e**), made fewer transitions between zones (**f**), and showed less exploratory activity (**g**, the number of grid lines crossed per second in the light zone (LZ, top) and dark zone (DZ, bottom)) than wild-type mice (black). **(h)** latency (s) to consume a novel food for α2^WT^ and α2^M/M^ mice in the hyponeophagia (HNP) test. Data are mean ± sem (**a-c**, n = 9; **d-g**, n = 8 and **h**, n = 9 (α2^WT^), with n = 15 (α2^M/M^)). ^#^p<0.05 and ^##^p<0.01 for the effect of genotype (α2^M/M^ vs wild-type), t test.

We then examined anxiety-based behaviour after pre-treating the mice with THDOC injected at doses known to be anxiolytic in both the EPM and LDB behavioural tests (*Wieland et al., 1991*; *Rodgers and Johnson, 1998*). In the EPM, 10 and 20 mg/kg THDOC proved anxiolytic in wild-type mice by increasing the relative number of open arm entries (*Figure 5a*) and the time spent on the open arms (*figure supplement 7*) compared to vehicle-injected control mice, without affecting their locomotor activity as determined from the number of closed and total arm entries (*Figure 5b – figure supplement 7*). By contrast, for α2^M/M^ mice on the EPM, 10 mg/kg THDOC did not reduce the aversion to the open arms, and only the 20 mg/kg THDOC dose was anxiolytic compared to control (*Figure 5a – figure supplement 7*). However, specifically for α2^M/M^ mice, THDOC injections unexpectedly reduced closed arm (*Figure 5b*) and total arm entries (*figure supplement 7*) suggesting a reduction in activity. Whilst this THDOC-induced effect might appear to confound interpretation, the relative open arm entries parameter inherently controls for the reduced activity. The knock-in mutation therefore displaced the anxiolytic dose-response relationship for THDOC towards higher doses, consistent with a reduced anxiolytic potency and/or efficacy of neurosteroids caused by the α2 subunit mutation.

**Figure 5.**
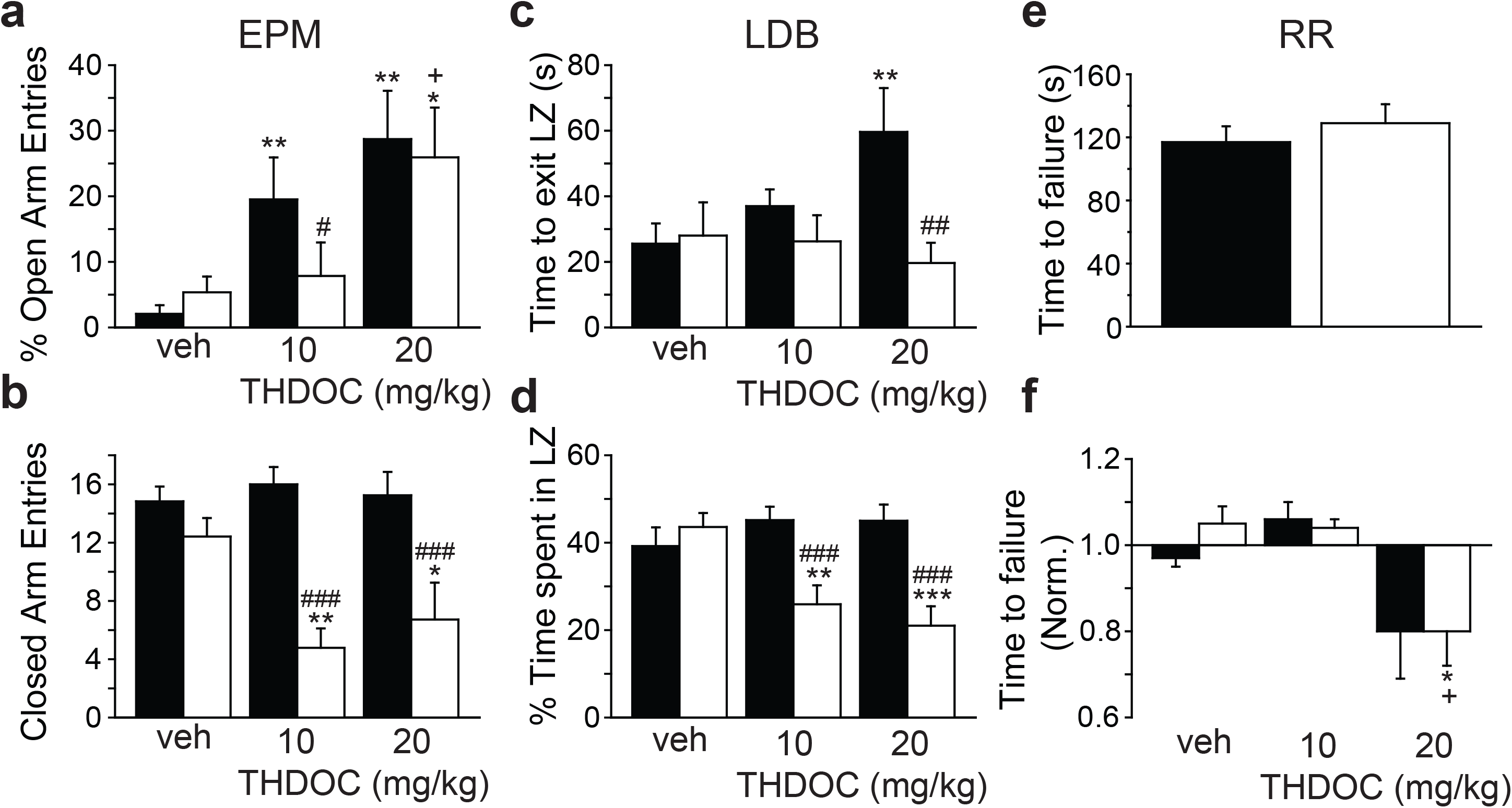
Anxiolytic effect of THDOC is impaired in α2^Q241M^ without affecting motor coordination. **(a and b)** Using the EPM, both 10 and 20 mg/kg THDOC were anxiolytic in wild-type mice (black), evidenced by increased open arm entries (**a**) without a change in locomotor activity (**b**) compared to vehicle (veh)-treated animals. Only 20 mg/kg THDOC was anxiolytic in α2^M/M^ mice (white) (**a**). **(c and d)** In the LDB, THDOC (20 mg/kg) was anxiolytic in wild-type mice (black) but not α2^M/M^ mice (white) as measured by the mean time to exit the light zone (LZ). (**c**) Injection of THDOC caused α2^M/M^ mice to spend less time in the light zone (**d**). **(e and f)** Motor coordination, examined by the time taken to fail the accelerating rotarod (RR) test; baseline coordination is no different between wild-type (black) and α2^M/M^ (white) mice (**e**). 20 mg/kg THDOC impairs performance on the rotarod in both genotypes relative to the untreated condition (**f**). Data are mean ± sem with * p<0.05, ** p<0.01, and *** p<0.001 for the effects of THDOC compared to vehicle; + p<0.05 for effect of 20 mg/kg THDOC vs. 10 mg/kg THDOC; # p<0.05, ## p<0.01 and ### p<0.001 for effect of genotype (i.e. α2^M/M^ vs. wild-type). Statistical tests used are: **a and b** (n = 7) and **c and d** (n = 9), ANOVA with the Benjamini-Hochberg adjustment; **e and f** (n = 7) comparisons used the Behrens-Fisher test.

For wild-type animals (α2^WT^) in the LDB, the anxiolytic profile of 20 mg/kg THDOC was evident from an increased time to exit the light zone (*Figure 5c*), and increased exploratory activity in both light and dark zones (*figure supplement 7*) compared to vehicle-injected control animals. In comparison, for α2^M/M^ mice, THDOC (10 and 20 mg/kg) caused no increase in the time to exit the light zone consistent with a lack of anxiolytic effect (*Figure 5c*). However, as with the EPM, THDOC seemingly reduced locomotor activity with reduced transfers between zones and less exploratory activity in either zone (*figure supplement 7*). THDOC also reduced the time spent in the light zone for the homozygote mice (*Figure 5d*), which could also reflect reduced activity following injection. However, THDOC-treated α2^M/M^ mice maintained their rapid egress from the light zone of the LDB (*Figure 5c*), suggesting the active avoidance response associated with the light zone clearly overcomes any THDOC activity-reducing effect. Overall, our observations with the light-dark box therefore concur with those from the elevated plus maze that the knock-in mutation α2^Q241M^ impairs the anxiolytic response to THDOC.

To determine whether the apparent reduced activity of α2^M/M^ mice represents a motor-impairing effect of injected THDOC, motor coordination was studied directly using the accelerating rotarod (RR) test. The Q241M mutation was not inherently motor-impairing as no basal genotypic differences were observed in initial performances on the rod (*Figure 5e*). After an initial training period (see Methods), mice were re-assessed following THDOC injection. No genotypic differences in performance were apparent at either THDOC dose tested (*Figure 5f*). Only at 20 mg/kg was THDOC motor-impairing (~ 20% decline in performance relative to pre-steroid condition), but this dose equally affected α2^WT^ and α2^M/M^ mice. Importantly, although 10 mg/kg THDOC has clear activity-reducing effects in α2^M/M^ mice in the EPM and LDB tests, it produced no motor impairment in the rotarod test (*Figure 5f*). Thus, the apparent activity-reducing effects of THDOC are not due to enhanced motor impairing effects in the knock-in mice.

### Depression and neurosteroid sensitivity of α2-GABA_A_Rs

The α2-GABA_A_R has also been closely associated with depression (*Smith and Rudolph, 2012*; *Mohler, 2012*). To examine the physiological contribution that neurosteroids may make to this behavioural phenotype, we first used the tail suspension test which equates a depressive phenotype to an increased amount of time spent in an immobile state following tail suspension (*Cryan et al., 2005*). Essentially, following tail suspension, the animals eventually adopt an immobile state following epochs of motor activity. These periods spent in an immobile posture (expressed as mean immobility time) for wild-type and α2^M/M^ mice were not significantly different, indicating that there are no basal anti-depressant effects of endogenous neurosteroids mediated via α2-GABA_A_Rs revealed by this test (*Figure 6a*). This lack of effect was corroborated by using another commonly applied test for assessing depression-type behaviour in rodents, the forced swim test (*Porsolt et al., 1977*). The initial tendency to escape the water is eventually replaced by immobility and this duration is considered to reflect a depressive phenotype. For both wild-type and knock-in mice the mean immobility times for this test were also similar (*Figure 6b*) again suggesting basal neurosteroid tone via α2-GABA_A_Rs plays little role in depression.

**Figure 6.**
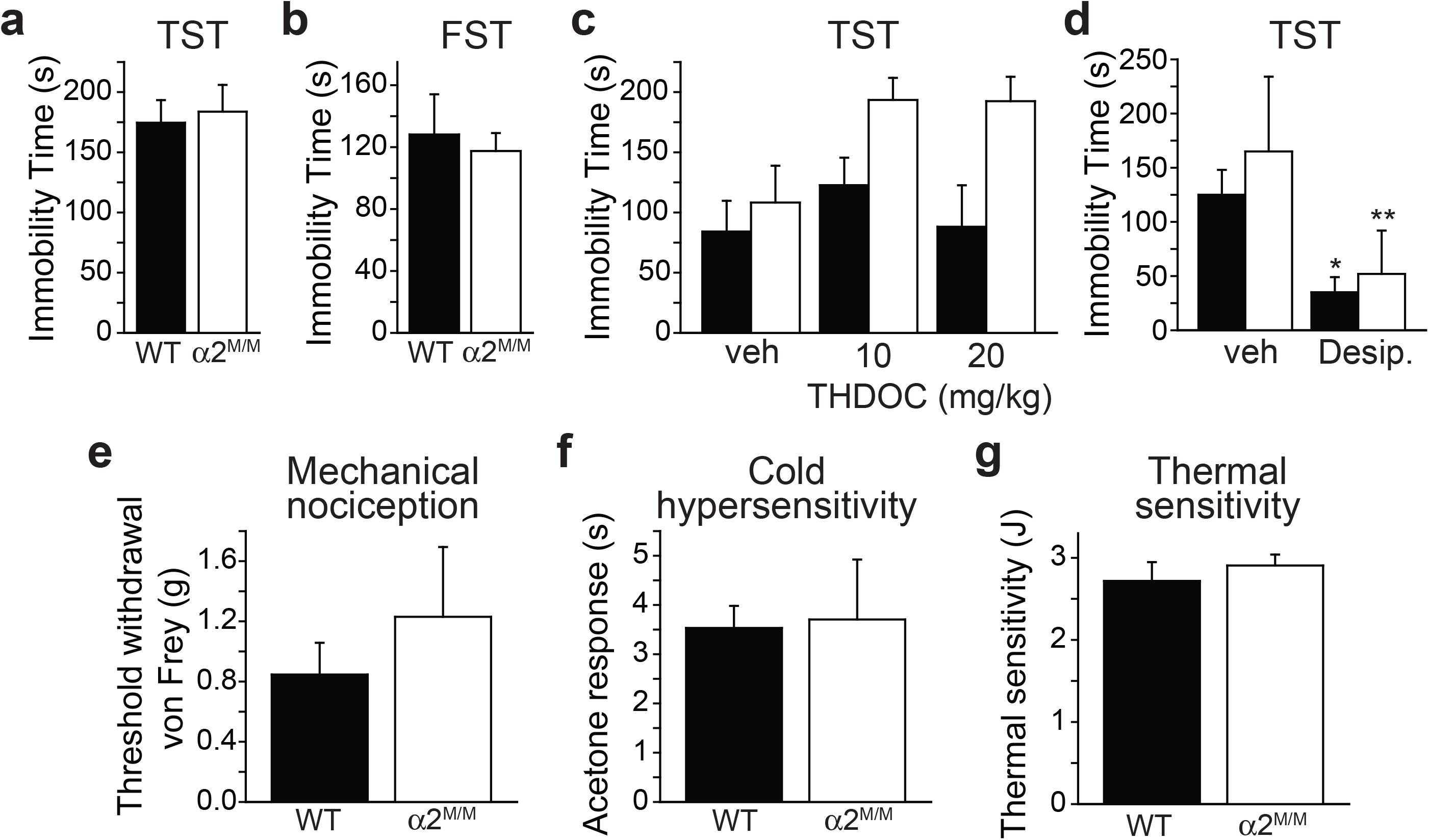
Basal neurosteroid tone on α2-GABA_A_Rs in acute depression and pain sensation. **(a)** Bar graph of the mean time spent immobile during the tail suspension test (TST) for wild-type (black) and α2^M/M^ (white) mice. There is no effect of genotype on basal depression-like behaviour (P>0.05; n = 8 per group). **(b)** Mean immobility time (s) in the forced swim test (FST) is the same for α2^WT^ and α2^M/M^ mice (n (α2^WT^) = 7; n (α2^M/M^) = 14; t-test, p=0.67. **(c)** THDOC is not antidepressant in wild-type or α2^M/M^ mice assessed by measuring the mean immobility time in the tail suspension test (P>0.05; n = 7 per group). **(d)** Bar graph of the mean time spent immobile following i/p administration of the anti-depressant, desipramine in the tail suspension test. Desipramine is effective in wild-type and α2M/M mice at 20 mg/kg. * - p<0.05, ** - p<0.01. For **a – d**, statistical comparisons were made using Bonferroni-corrected, Fisher’s least significant difference test. P values are for all comparisons, n = 7 - 14 per group. **(e-g)** Bar graphs measuring hind-limb withdrawal reflexes for mechanical nociception (**e**, using von Frey filaments); cold hypersensitivity (**f**, using acetone droplets), and thermal sensitivity (**g**, using laser light). Units in plots are: g = grams, s = seconds, J = Joules. No differences were observed between α2^WT^ and α2^M/M^ mice (P> 0.05; n = 8). Statistical comparisons were made using the t-test.

We next explored whether injected neurosteroids could have an anti-depressant effect by using the tail suspension test. However, injecting THDOC at 5 or 10 mg/kg into both genotypes did not affect the immobility times for either α2^WT^ or α2^M/M^ mice, when compared to vehicle-injected controls (*Figure 6c*). As a control for the tail suspension test being indicative of depression, we also examined the effect of the anti-depressant, desipramine. Using an anti-depressant dose of 20 mg/kg (Ref (*Cryan et al., 2005*)), desipramine significantly reduced the mean times spent in an immobile posture for both wild-type and α2^M/M^ mice when compared to vehicle-injected controls (*Figure 6d*).

### Pain sensation and neurosteroid sensitivity of α2-GABA_A_Rs

GABA_A_Rs are implicated in the process of pain sensation, particularly those containing α2 subunits, and analgesia can also be attained by targeting receptors that contain α2 or α3 subunits (*Zeilhofer et al., 2009*; *Zeilhofer et al., 2012*; *Knabl et al., 2008*). In addition, inflammatory pain can lead to enhanced spinal inhibition following an upregulated production of neurosteroids (*Poisbeau et al., 2005*). Accordingly, we examined whether a basal neurosteroid tone on α2-GABA_A_Rs was important for normal pain sensation by measuring the hindlimb withdrawal reflex, following exposure to a range of nociceptive modalities, including: mechanical nociception (using von Frey filaments); cold hypersensitivity (using droplets of acetone applied to the skin); and thermal sensitivity (with a focused laser light applied to the skin). For all three assessments of pain, the wild-type mice showed equal sensitivity compared with α2^M/M^ mice, indicating that a basal neurosteroid tone acting via α2-GABA_A_Rs was unlikely to be important for innate pain sensation (*Figure 6e-g*).

## Discussion

There are many putative physiological roles that are attributed to the neurosteroids in the brain ranging from anxiolytic, anti-depressant, anti-convulsant, and anti-stress, to roles in cognition, learning and memory (*Reddy, 2010*). Of these, we have established here that their innate ability to act as anxiolytics (*Purdy et al., 1991*; *Barbaccia et al., 1998*; *Wieland et al., 1991*; *Crawley et al., 1986*) principally depends upon potentiation of GABA activation at α2 subunit-containing GABA_A_Rs. The introduction of a single point mutation into the α2 receptor in a mouse line, replacing H-bonding glutamine with hydrophobic methionine (Q241M), was singularly sufficient to ablate neurosteroid activity via these receptors. This residue switch demonstrated unequivocally that GABA_A_Rs are natural targets for endogenous neurosteroids. Moreover, at basal levels, neurosteroids modulate synaptic inhibition in neural circuits that are closely associated with anxiety in the hippocampus and the medial prefrontal cortex. This important role resonates with the reported ability of other positive allosteric modulators, namely subtype-selective benzodiazepines to target α2 and α3 subunit-containing GABA_A_Rs and exhibit anxiolytic profiles (*Low et al., 2000*; *Dias et al., 2005*).

We used patch clamp electrophysiology to reveal the basis for neurosteroid modulation of GABA-mediated inhibition at α2 subunit-containing GABA_A_Rs in hippocampal and cortical slices, which revealed some unexpected facets. Firstly it was clear on comparing recordings from neurons in α2^WT^ and α2^M/M^ slices from both brain regions that there is an endogenous neurosteroid tone that modulates GABA transmission. By using the 5α-reductase inhibitor finasteride to remove neurosteroids from hippocampal slices, the effect of the α2^Q241M^ mutation on DGGC sIPSC decay times was abolished. A similar tone has been noted previously in the neocortex, revealed by inhibiting neurosteroid synthesis (*Puia et al., 2003*), but it has not been resolved at the level of individual GABA_A_R isoforms.

Whilst α2-GABA_A_Rs are considered to be representative of classical synaptic-type GABA_A_ receptors, the extent of inhibition mediated by this receptor subtype in hippocampal CA1 and layer II/II mPFC pyramidal neurons was far larger than expected. By using the loss of THDOC modulation in the α2^Q214M^ line as an index of α2 subunit involvement, GABA_A_Rs with this subunit seemingly accounted for a high proportion of inhibition in both cell types since THDOC prolonged the decay time of sIPSCs by ~25-30% for wild-type neurons compared to only ~10-15% for α2^M/M^ counterparts. Although neurosteroid modulation of GABA synaptic transmission is reduced in both α2^M/M^ pyramidal neuron populations, it is not blocked, and presumably other GABA_A_R isoforms, with intact neurosteroid sensitivity, are available to mediate the sIPSC decay. Interestingly, in dentate gyrus granule cells, α2^M/M^ is sufficient to abolish neurosteroid modulation of sIPSC decays suggesting an even larger role for α2 GABA_A_Rs in mediating synaptic inhibition in this area of the hippocampus. It is unlikely that GABA_A_R subunit compensation is affecting synaptic inhibition since our western blot and immunohistochemical data indicated no changes to the expression patterns of the major α subunits. Moreover, the benzodiazepine diazepam equally prolonged the sIPSC decays for α2^WT^ and α2^M/M^ in dentate gyrus neurons. Also, the amplitudes of the sIPSCs remained largely unaffected by α2^M/M^ indicating there are no major changes to the levels of expressed synaptic GABA_A_Rs at the cell surface. Taken overall, these data suggest that α2 subunit GABA_A_Rs play a key role in synaptic transmission and in the modulatory effects of the neurosteroids.

An unexpected finding from examining GABA inhibition in the α2 knock-in mouse line was the clear evidence that α2 subunit-containing receptors can mediate tonic inhibition in the dentate gyrus, most likely via receptors of an as-yet unreported subunit composition. Addition of THDOC caused the predicted increase in tonic current in dentate neurons and this was remarkably reduced by ~74 % in α2^M/M^ neurons implying an important role for α2 subunits in the neurosteroid modulation of tonic GABA inhibition. By comparison, tonic current was largely insensitive to diazepam, eliminating the likelihood that extrasynaptic GABA_A_Rs containing γ2 subunits are supporting the tonic current with α2 subunits. What then is the likely subunit composition of the α2 extrasynaptic GABA_A_Rs? We know that GABA_A_Rs absolutely require co-assembly of a β subunit for efficient cell surface expression (*Connolly et al., 1996a*; *Connolly et al., 1996b*) and if the γ2 subunit is not required following the insensitivity of the tonic current to diazepam, then α2 subunit receptors mediating tonic inhibition are most likely to be perisynaptic or extrasynaptic receptors comprising a combination of α2β or α2βδ subunits – both of which are novel isoforms of the GABA_A_R (*Olsen and Sieghart, 2009*).

The importance of the neurosteroid tone via α2-GABA_A_Rs became evident from the behavioural tests. Three tests for anxiety suggested removing the basal endogenous neurosteroid modulation in α2^M/M^ mice caused an anxiogenic phenotype. This was manifest by α2^M/M^ mice spending less time on the open arms of the elevated plus maze and in the light dark box being quicker to leave, spending less time in, and making fewer returns to, the light zone. The reduced activity noted in the dark zone is also in accord with a heightened state of anxiety. In addition, the increased latency to sample novel food in the hyponeophagia test for α2^M/M^ is again indicative of increased basal anxiety.

By raising neurosteroid levels using THDOC injections into α2^WT^ and α2^M/M^ mice, divergent effects were apparent. For α2^WT^ using the EPM, clear anxiolytic effects were manifest by increased open arm entries and increased time spent on the open arms without any activity changes. Similarly in the LDB, there was no inclination to exit the light zone and mice also showed increased general exploration. However, for the α2^M/M^ mouse, it was apparent that THDOC seemingly reduced locomotor activity in both the EPM and LDB. This reduction in activity is not caused by a THDOC-induced motor impairment in the knock in mice since the rotorod test revealed no differences in motor performance for α2^WT^ and α2^M/M^ even at the highest doses of THDOC. Whilst the THDOC-induced reduction in activity makes clear interpretation of anxiety-motivated behaviour more difficult, there is sufficient compelling evidence that the anxiolytic effect of THDOC is compromised in the α2^M/M^ mice. In the EPM the proportion of open arm entries was less sensitive to being increased by THDOC, and, in the LDB, THDOC was clearly ineffective at increasing the time taken to exit the light zone. Of importance, both these parameters should be uncompromised by any apparent reduction in activity. Hence, our data indicates that the anxiolytic effects mediated by both basal and elevated levels of neurosteroids are significantly diminished by removing modulation at α2-GABA_A_Rs.

It is frequently noted that anxiety and depression are co-morbid (*Mohler, 2012*). These disease phenotypes are often the result of repeated stress episodes, which can affect nervous system activity by causing variations in neurosteroid levels (*Gunn et al., 2011*; *Maguire and Mody, 2007*; *Sarkar et al., 2011*), the results of which are manifest by signalling via GABA_A_Rs and consequently altered behaviour.

There is growing evidence to support a role for α2 and α3 subunit-containing GABA_A_Rs in depression, from the use of subtype-selective modulators (*Mohler, 2012*). However, from our study, at least in terms of basal neurosteroid tone, signalling via α2-GABA_A_Rs does not perform a physiological role in depression ascertained with the tail suspension text and forced swim test. Moreover in the former test, a classical anti-depressant, desipramine, is equally effective at reducing depressive-like behaviour for both the wild-type and α2^M/M^ animals.

GABA_A_Rs have also been reported to exert some basal control over neuronal responses to pain (*Reeve et al., 1998*). Furthermore, a reduction in GABA_A_R mediated synaptic inhibition is important in the dorsal horn of the spinal cord for the development of pain, such that GABA_A_Rs are considered to be therapeutic targets for analgesic agents (*Knabl et al., 2008*; *Zeilhofer et al., 2012*). In particular, targeting α2- and α3-GABA_A_Rs with selective benzodiazepine agents has provided potentially useful analgesics for treating inflammatory and neuropathic pain (*Knabl et al., 2008*; *Zeilhofer et al., 2009*). Our examination of basal neurosteroid tone acting via α2-GABA_A_Rs in mediating pain sensation, as manifest by measurements of mechanical, cold and thermal sensitivities, indicated a minimal, if any, role for basal neurosteroid signalling via these GABA_A_Rs.

Overall, our knock-in animal model, which lacks neurosteroid potentiation at α2-GABA_A_Rs, reveals that neurosteroids are physiologically active in the brain. It provides the first direct evidence that they can act as endogenous anxiolytics in a remarkably specific, GABA_A_R isoform selective manner, with little or no impact on depression and pain sensation, both of which have previously been associated with α2-GABA_A_Rs. These findings form a platform for the development of therapeutic agents for the specific treatment of anxiety, and stress-related mood disorders, relatively unencumbered by debilitating side effects which blight so many current clinical therapies. Furthermore, on the basis of ablating neurosteroid action via α2 GABA_A_Rs, we predict that other important physiological and pathophysiological actions of neurosteroids are likely to co-associate with specific GABA_A_R isoforms and neural circuits.

## Materials and Methods Animals

All procedures were performed in accordance with the Animals (Scientific Procedures) Act 1986. The α2^Q241M^ knock-in mice were generated as described in the Supplementary Information, and maintained on a C57BL/6J background. Given that steroid hormones of the oestrus cycle have a strong influence on brain neurosteroid levels in female mice (*Corpechot et al., 1997*), our experiments used only males.

A targeting vector containing exon 8 of the GABA_A_R α2 subunit gene with base pair changes (CAA to ATG) for the point mutation Q241M (*figure supplement 1*) was made utilising an RP23 Bacterial Artificial Chromosome library clone derived from a C57BL/6J mouse. Homologous recombination (GenOway, Lyon, France) was achieved by introducing the targeting vector into 129Sv/Pas embryonic stem (ES) cells using electroporation. PCR and Southern blotting techniques identified ES cell lines that incorporated the Q241M mutation (the mutagenesis silently created an Nco-I restriction site, allowing identification of homologous recombinants by the change in size of Nco-I digestion products – see *Figure 1a*). These ES cell lines were used to generate chimeric mice expressing α2^Q241M^. Germline-transmitted pups from a chimera x C57BL/6J cross (F0 generation) were bred with C57BL/6J Cre-recombinase-expressing mice to remove the neomycin resistance cassette (F1 generation – see *figure supplement 1*). Further backcrosses were performed between heterozygote α2^Q241M^ mice and C57BL/6J mice. The mice used in the experiments described here are from generations F4 - F6. Animals were maintained as heterozygous pairs.

### Antibodies

For Western blotting, the following primary antibodies were used: mouse anti-α1-GABA_A_R (1:1000 dilution, Neuromab), guinea-pig anti-α2-GABA_A_R (1:1000, Synaptic Systems), rabbit anti-α2-GABA_A_R (1:500, from Werner Sieghart, University of Vienna), rabbit anti-α3-GABA_A_R (1:1000, Alomone Labs), rabbit anti-α4-GABA_A_R (from Werner Sieghart), mouse anti-β-tubulin (Sigma-Aldrich). The following horseradish-peroxidase-conjugated secondary antibodies (IgG(H&L)) were used: goat anti-rabbit (1:10000, Rockland Immunochemicals), goat anti-mouse (1:10000, Rockland Immunochemicals), donkey anti-guinea-pig (1:1000, Jackson Immunoresearch Laboratories).

For immunohistochemistry, the following primary antibodies were used: rabbit anti-α1-GABA_A_R (1:20000, from Jean-Marc Fritschy, University of Zurich), guinea-pig anti-α2-GABA_A_R (1:1000, from Jean-Marc Fritschy), rabbit anti-α3-GABA_A_R (1:1000, Alomone Labs), rabbit anti-α4-GABA_A_R (1:500, Werner Sieghart), rabbit anti-α5-GABA_A_R (1:500, Werner Sieghart). Alexa Fluor 555-conjugated goat anti-rabbit and goat anti-guinea pig (IgG (H&L)) were purchased from Invitrogen Ltd and used at 1:2000.

### Western blotting

Hippocampi and cortices from postnatal age P18 - 30 mice were lysed to isolate total protein. Equal amounts of total protein were subjected to Western blotting to assess the expression of each GABA_A_R α subunit, using β-tubulin as a loading control. Total protein was isolated from each brain area by homogenisation in ice-cold RIPA buffer, followed by cell disruption with repeated freeze-thaw cycles. Equal amounts of total protein were subjected to sodium-dodecyl-sulphate polyacrylamide gel electrophoresis and transferred to nitrocellulose membranes. Blotting was performed using 4% w/v milk in 0.1% v/v TWEEN-supplemented phosphate buffered saline (PBS), with blocking for 1 h at room temperature (RT), exposure to primary antibody overnight at 4°C and secondary antibody for 2 h at RT. Following incubation with each antibody, membranes were washed 3x (20 min, RT). Blots were finally rinsed with PBS and developed using chemiluminescent substrate and visualised with an ImageQuant LAS4000 imager. Images were quantified using the Western blot plug-in on ImageJ software (Version 1.44p, National Institutes of Health, USA). After blotting for the relevant GABA_A_R α subunit, blots were subjected to a mild stripping procedure (10 - 20 min incubation in a buffer comprising 200 mM glycine, 0.1% w/v SDS, 1% v/v TWEEN-20, pH 2.2), washed and re-blotted for quantifying β-tubulin expression. Three samples of each genotype were run on each blot, and three blots were performed (i.e. 9 each of wild-type, α2^Q/M^ and α2^M/M^ samples in total). The density of each α subunit band was first normalised to its corresponding β-tubulin band, then signals within each Western blot were normalised to the average signal of the wild-type bands in that blot.

### Immunofluorescence

Cardiac perfusion and tissue sectioning were performed as previously described (*Tochiki et al., 2012*). Slices were permeabilised/blocked for 1h at RT in permeabilisation/blocking solution (0.5% v/v bovine serum albumin, 2% v/v normal goat serum, 0.2% v/v triton-x-100 in PBS). Primary (overnight, 4°C) and secondary (2 h, RT) antibody incubations were performed in the same permeabilisation/blocking solution. After each antibody incubation, slices were washed 4x in PBS (15 min, RT). Controls for non-specific labelling by secondary antibodies were performed by staining as described, but omitting primary antibody. Images were acquired using a Zeiss LSM510 confocal microscope (Carl Zeiss) with a 63x immersion objective. Immunofluorescent staining assessed the expression of α1 - α5 subunits in coronal hippocampal slices (40 μm) from P42 - P70 mice. The α6 subunit isoform was not studied because of its restricted spatial expression to granule cells of the cerebellum and the cochlear nucleus (*Laurie et al., 1992a*; *Pirker et al., 2000*). For each subunit and cell-type, three slices were imaged per animal, capturing data as z-stacks, with images in each plane acquired as a mean of 8 scans in 8 bits. Blinded image analyses were performed using ImageJ (Version 1.44p) to determine the mean fluorescence intensity for separate regions of interest for the dendrites and cell bodies. In both cases, to ensure that the value is representative of the staining throughout the brain slice, three readings were taken per image: one from the top, one central and one at the bottom of the z-stack; the average of these values was taken to represent the expression of the subunit in that slice. Results are normalised to the mean fluorescence intensity observed in wild-type slices.

### Electrophysiology Recombinant GABA_A_Rs

Whole-cell currents were recorded from transfected HEK-293 cells 48 hr after transfection. Cells were voltage clamp at −30 mV with series resistances less than 10 MΩ. Borosilicate glass patch electrodes (resistances of 3–5 MΩ) were filled with an internal solution containing (mM): 120 CsCl, 1 MgCl_2_, 11 EGTA, 30 KOH, 10 HEPES, 1 CaCl_2_, and 2 K_2_ATP; pH – 7.2. HEK-293 cells were superfused with a solution containing (mM): 140 NaCl, 4.7 KCl, 1.2 MgCl_2_, 2.52 CaCl_2_, 11 Glucose, and 5 HEPES; pH 7.4. Membrane currents were filtered at 5 kHz (−3 dB, 6th pole Bessel, 36 dB per octave).

Concentration response relationships for GABA-activated membrane currents, in the presence and absence of THDOC, were compiled by measuring the current (I) at each GABA concentration and normalizing these currents to the maximal GABA current (I_max_). The concentration response relationship was fitted with the Hill equation:

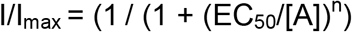

where A is the concentration of GABA, EC_50_ is the concentration of GABA causing 50% of the maximal GABA current, and n is the slope. All concentration response data were curve fitted in Origin (Ver 6).

### Brain slices

GABA-mediated membrane currents were recorded from P18 - P30 coronal (350 μm) hippocampal slices. Slices were prepared in ice-cold artificial cerebrospinal fluid (aCSF) comprising (mM): 130 K-gluconate, 15 KCl, 0.05 EGTA, 20 HEPES, 4 Na-pyruvate, 25 glucose and 2 kynurenic acid, pH 7.4. Over 1 h, slices were incubated at 37°C, whilst the solution was exchanged for electrophysiology to aCSF (mM): 125 NaCl, 2.5 KCl, 1.25 Na_2_H_2_PO_4_, 26 NaHCO_3_, 2 CaCl_2_, 1 MgCl_2_, 25 glucose. All aCSF solutions were continuously bubbled with 95%O_2_/5%CO_2_ (BOC Healthcare). Whole-cell recordings were undertaken at RT from single neurons located using infra-red optics and patch pipettes (3 - 5 MΩ) filled with an internal solution containing (mM): 140 CsCl, 2 NaCl, 10 HEPES, 5 EGTA, 2 MgCl_2_, 0.5 CaCl_2_, 2 Na-ATP, and 2 QX-314 bromide. Slices were continuously perfused with recording aCSF supplemented with kynurenic acid, to isolate GABAergic synaptic events. THDOC and diazepam were prepared as stock solutions (10 mM and 100 mM, respectively) in DMSO, and diluted to the appropriate final concentration in recording aCSF. Drugs were allowed to equilibrate in the bath for at least 5 min; control or ‘mock’ recordings were performed with the DMSO vehicle. Experiments were terminated if series resistance changed by more than 20%. Properties of individual sIPSCs were assessed by fitting at least 100 isolated events using MiniAnalysis software (Synaptosoft Inc., New Jersey, USA) to define the average synaptic event rise time, amplitude and decay times. Mono- and bi-exponentially decaying events were analysed together by transforming to weighted decay time, τ_w_, according to the equation

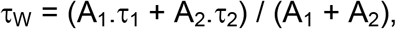

where τ_1_ and τ_2_ are exponential decay time constants, and A_1_ and A_2_ are the relative amplitude contributions of τ_1_ and τ_2_, respectively (for mono-exponential events: A_2_ = 0). The frequency of sIPSCs was measured from the inter-event interval for all events within a 60 s recording period. Results are expressed as a percentage change for the test epoch relative to the control epoch. Tonic currents were measured by applying 20 μM bicuculline which reduced the holding current. These were analysed with WinEDR (version 3.1; J Dempster, University of Strathclyde, UK). Effects of THDOC or diazepam on tonic current were measured from absolute changes relative to controls. To examine sIPSC kinetics in the absence of endogenous neurosteroids, biosynthesis was blocked by supplementing aCSF solutions with 50 μM finasteride. Slices were incubated in this solution for at least 6 h before recording, to allow for the clearance of neurosteroids already present in the slice before blocking the 5α-reductase enzyme.

### Behaviour

Experimentally naïve mice (P42 - P108) were used in single behavioural tests. Each test used a separate cohort of mice and typically each cohort contained 7 – 10 mice. When used, THDOC was dissolved in 0.9% w/v NaCl with 25% w/v 2-hydroxypropyl-β-cyclodextrin and administered intraperitoneally at 10 ml per kg body weight 15 min before the start of the test. Note that for the light dark box and elevated plus maze testing, animals were tested in pairs (one α2^M/M^ mouse and one wild-type), and the drug dose received and the order in which mice were tested (wild-type first vs. α2^M/M^ first) was randomised. Within the rotarod test, the drug dose administered to each mouse was pre-determined by a counterbalanced randomisation procedure. Scoring was performed blind to both the genotype and drug administered. Blinding used a random number based method independently applied to the investigator.

### Elevated Plus Maze

The EPM apparatus was as described previously (*Lister, 1987*), except that the walls on the closed arms were black. The maze was elevated 500 mm above the floor of the experimental room, and the illumination across the arms was uniform and low intensity (approximately 25 lux, equivalent to 25 lumen per m^2^). Mice were placed in the centre of the maze, initially facing one of the closed arms, and movements within the maze over a 5 min trial were monitored manually with a single scorer blinded to genotype with a digital camcorder mounted 1.5 m directly above the maze. The percentage time in an arm is expressed as (time in arm / total time of experiment) x 100. Entry into an arm was defined by all four paws crossing the arm threshold.

### Light-Dark Box

LDB dimensions were 20 x 45 cm, which were divided into a light zone (two-thirds total area, illumination 850 lux) and a dark zone (one-third total area, 50 lux) connected by a small doorway. The floors of each zone were marked with a grid to allow scoring of exploratory activity as lines crossed per unit time. A mouse was placed initially in the light zone, facing away from the doorway, and was then allowed to freely explore the box for 10 min. Mouse movements were monitored using video recording and exploratory activity scored blind to genotype.

### Hyponeophagia

The hyponeophagia test was performed as previously described (*Deacon, 2011*). Briefly, mice were deprived of food for 12-16 hrs prior to testing (1g lab food per mouse). Prior to testing, mice were moved to individual cages for 30 min to avoid social transmission of any preference for the novel food. Animals were positioned facing away from the food placed on a small platform containing a well filled with ~1 ml condensed milk (Nestle carnation; diluted 50:50 with water). Between trials the setup was cleaned with 70% v/v ethanol. The time taken to consume the milk (>2s continuous drinking) was measured by an observer blinded to the genotype of the animals. If an animal did not consume any milk within 3 min the trial was terminated.

### Tail Suspension test

For the tail suspension test, mice were secured to vertical rods using hypoallergenic, inert tape applied to the tails, in 5 discrete chambers. The test duration was 6 mins and the total amount of time they spent in an immobile state was recorded (mean immobility time) once this had occurred for more than 20 s.

### Forced Swim test

The apparatus consisted of a large beaker (diameter: 20 cm; height: 35 cm) half-filled with water at 22-23°C. Animals were transferred to individual cages inside the experimental room 45 min before testing. During the test, animals were carefully lowered into the beaker for 6 min while their behaviour was monitored with a webcam (PS3 eye camera). After the trial was complete each animal was dried with a towel before being returned to its home cage. The time spent immobile was scored from the video files for the last 4 min of each trial by an observer blinded to the genotype of the mouse according to criteria previously described (*Can et al., 2012*).

### Rotarod

Mice were placed on a five-station rotarod treadmill apparatus (Med Associates Inc., Vermont, USA) with the rod rotating initially at 4 r.p.m. The rod accelerated uniformly to a final speed of 40 r.p.m. over 5 min. Scoring assessed the time at which a mouse failed the test, either by falling from the rod, or by passively rotating with the rod. If a mouse completed the full test, the ‘time to failure’ was simply set as the duration of the test (300 s). The performance of each mouse on their first encounter with the rod was used as a measure of the baseline motor coordination. Each mouse was subjected to further training (5 trials per day for 4 consecutive days), such that a consistent performance was attained before the effect of drug was assessed. Any impairment of motor coordination caused by THDOC was determined by expressing this performance relative to their average performance on the rod on the preceding drug-free training day.

### Pain

Mechanical sensitivity – Mice were acclimatised to the behavioural suite for 1 hr prior to testing. Animals were then placed in a chamber containing a wire mesh floor and allowed another 15 min to settle. Mechanical sensitivity was then assessed using von Frey filaments (Touch-Test, North Coast Medical, USA) applied to the hindlimb until they buckled for 5 - 6 s. Any withdrawal, lifting, flinching and shaking were defined as a positive response. We determined the threshold for 50% withdrawal using the up-down method as previously described (*Chaplan et al., 1994*) with von Frey filaments delivering forces of 0.07 to 0.4 g.

Cold sensitivity – This was assessed by applying a drop of acetone using a 0.5 ml syringe to the upper surface of one of the hind paws. Acetone was applied 5 times, with at least 3 min recovery between applications. The number of withdrawals out of 5 was recorded. Licking, flinching and shaking were considered as positive responses.

Thermal sensitivity – The withdrawal reflex was measured by focussing an infrared neodymium-doped yttrium aluminium perovskite (Nd:YAP; λ 1.34 μm) laser onto the upper surface of one of the hindlimbs (*Patel et al., 2013*) and then increasing laser power, over a range of 1 – 3.5 J, until a thermal threshold was observed, ascertained by hindlimb withdrawal. A thermal cut-off was set at 3.5 J to avoid tissue damage.

### Statistics

Parametric comparisons were performed for normally distributed, equal variance data (analysis of variance (ANOVA) and t-test). Where data did not satisfy these criteria, they were first transformed (e.g. logarithmic transform), and the criteria reassessed before proceeding with any parametric comparison. ANOVA comparisons (with the Benjamini-Hochberg adjustment) were performed using InVivoStat (British Association for Psychopharmacology, Cambridge, UK), and used to report the Fisher’s least-significant difference (LSD) p-values from planned individual pairwise comparisons. Where data transformation was unable to satisfy the criteria for parametric comparison, non-parametric analysis was used, e.g., Kruskal-Wallis test. Where appropriate, and only if the overall Kruskal-Wallis test indicated a significant variation between discrete groups, individual pairwise comparisons were performed using the Mann-Whitney test.

## Acknowledgements

This work was supported by the Medical Research Council UK

## Author Information

The authors declare no competing financial interests. Correspondence and request for materials should be addressed to TGS.

## Acknowledgements

We thank Clare Stanford for considerable help and advice with the behavioural experiments, and Stephen Hunt for the loan of elevated plus maze and rotarod equipment. This work was supported by the MRC and Sinergia.

The following figure supplements are available for figure 1:

**Figure supplement 1.** Targeting GABA_A_R α2 subunits to create a neurosteroid binding site knock-in mouse line.

**Figure supplement 2.** Immunofluorescent staining in CA1 of the hippocampus

**Figure supplement 3.** Immunofluorescent staining in the dentate gyrus

**Figure supplement 4.** Unaltered GABA_A_R α1 - α5 subunit immunofluorescence in the α2Q241M knock-in.

The following figure supplements are available for figure 2:

**Figure supplement 5.** Synaptic currents recorded from mPFC pyramidal neurons in α2Q241M knock-in slices.

**Table supplement 1.** Properties of sIPSCs across genotypes

**Table supplement 2.** THDOC-induced changes in CA1 PN sIPSCs

**Table supplement 3.** Drug-induced changes in DGGC sIPSCs

**Table supplement 4.** THDOC-induced changes in layer II/III mPFC PN sIPSCs

The following figure supplements are available for figure 3:

**Figure supplement 6.** THDOC modulation of recombinant synaptic and extrasynaptic-type GABA_A_Rs

**Table supplement 5.** Tonic currents in DGGCs

**Table supplement 6.** Tonic currents in layer II/III mPFC PNs

**Table supplement 7.** THDOC modulation of recombinant synaptic- and extrasynaptic-type GABA_A_Rs

The following figure supplements are available for figure 4:

**Figure supplement 7.** Additional data from EPM and LDB tests

The following figure supplements are available for figure 5:

**Figure supplement 7.** Additional data from EPM and LDB tests

## References

Barbaccia ML, Colombo G, Affricano D, Carai MA, Vacca G, Melis S, Purdy RH, Gessa GL. 2002. GABA_B_ receptor-mediated increase of neurosteroids by γ-hydroxybutyric acid. Neuropharmacology. 42: 782–791.

Barbaccia ML, Concas A, Serra M, Biggio G. 1998. Stress and neurosteroids in adult and aged rats. Exp Gerontol. 33: 697–712.

Belelli D and Herd MB. 2003. The contraceptive agent Provera enhances GABA_A_ receptor-mediated inhibitory neurotransmission in the rat hippocampus: evidence for endogenous neurosteroids? J.Neurosci. 23: 10013–10020.

Belelli D, Herd MB, Mitchell EA, Peden DR, Vardy AW, Gentet L, Lambert JJ. 2006. Neuroactive steroids and inhibitory neurotransmission: mechanisms of action and physiological relevance. Neuroscience. 138: 821–829.

Can A, Dao DT, Arad M, Terrillion CE, Piantadosi SC, Gould TD. 2012. The Mouse Forced Swim Test. Journal of Visualized Experiments : JoVE. 3638.

Chaplan SR, Bach FW, Pogrel JW, Chung JM, Yaksh TL. 1994. Quantitative assessment of tactile allodynia in the rat paw. J Neurosci Methods. 53: 55–63.

Concas A, Mostallino MC, Porcu P, Follesa P, Barbaccia ML, Trabucchi M, Purdy RH, Grisenti P, Biggio G. 1998. Role of brain allopregnanolone in the plasticity of γ-aminobutyric acid type A receptor in rat brain during pregnancy and after delivery. Proc.Natl.Acad.Sci U.S.A. 95: 13284–13289.

Connolly CN, Krishek BJ, McDonald BJ, Smart TG, Moss SJ. 1996a. Assembly and cell surface expression of heteromeric and homomeric gamma-aminobutyric acid type A receptors. J.Biol.Chem. 271: 89–96.

Connolly CN, Wooltorton JR, Smart TG, Moss SJ. 1996b. Subcellular localization of gamma-aminobutyric acid type A receptors is determined by receptor beta subunits. Proc.Natl.Acad.Sci.U.S.A. 93: 9899–9904.

Corpechot C, Collins BE, Carey MP, Tsouros A, Robel P, Fry JP. 1997. Brain neurosteroids during the mouse oestrous cycle. Brain Res. 766: 276–280.

Crawley JN, Glowa JR, Majewska MD, Paul SM. 1986. Anxiolytic activity of an endogenous adrenal steroid. Brain Res. 398: 382–385.

Cryan JF, Mombereau C, Vassout A. 2005. The tail suspension test as a model for assessing antidepressant activity: Review of pharmacological and genetic studies in mice. Neuroscience & Biobehavioral Reviews. 29: 571–625.

Deacon RM. 2011. Hyponeophagia: A Measure of Anxiety in the Mouse. Journal of Visualized Experiments : JoVE. 2613.

Dias R, Sheppard WF, Fradley RL, Garrett EM, Stanley JL, Tye SJ, Goodacre S, Lincoln RJ, Cook SM, Conley R, Hallett D, Humphries AC, Thompson SA, Wafford KA, Street LJ, Castro JL, Whiting PJ, Rosahl TW, Atack JR, McKernan RM, Dawson GR, Reynolds DS. 2005. Evidence for a significant role of α3-containing GABA_A_ receptors in mediating the anxiolytic effects of benzodiazepines. J.Neurosci. 25: 10682–10688.

Dixon CI, Rosahl TW, Stephens DN. 2008. Targeted deletion of the *GABRA2* gene encoding α2-subunits of GABA_A_ receptors facilitates performance of a conditioned emotional response, and abolishes anxiolytic effects of benzodiazepines and barbiturates. Pharmacol Biochem Behav. 90: 1–8.

Engin E, Smith KS, Gao Y, Nagy D, Foster RA, Tsvetkov E, Keist R, Crestani F, Fritschy JM, Bolshakov VY, Hajos M, Heldt SA, Rudolph U. 2016. Modulation of anxiety and fear via distinct intrahippocampal circuits. Elife. 5: doi.org/10.7554/eLife.14120.001.

Engin E and Treit D. 2007. The anxiolytic-like effects of allopregnanolone vary as a function of intracerebral microinfusion site: the amygdala, medial prefrontal cortex, or hippocampus. Behav.Pharmacol. 18: 461–470.

Farrant M and Nusser Z. 2005. Variations on an inhibitory theme: phasic and tonic activation of GABA_A_ receptors. Nat.Rev.Neurosci. 6: 215–229.

Gunn BG, Brown AR, Lambert JJ, Belelli D. 2011. Neurosteroids and GABA_A_ Receptor Interactions: A Focus on Stress. Front Neurosci. 5: 131.

Harney SC, Frenguelli BG, Lambert JJ. 2003. Phosphorylation influences neurosteroid modulation of synaptic GABA_A_ receptors in rat CA1 and dentate gyrus neurones. Neuropharmacology. 45: 873–883.

Helms CM, Rossi DJ, Grant KA. 2012. Neurosteroid influences on sensitivity to ethanol. Front Endocrinol.(Lausanne). 3: 10.

Hirani K, Khisti RT, Chopde CT. 2002. Behavioral action of ethanol in Porsolt’s forced swim test: modulation by 3 alpha-hydroxy-5 alpha-pregnan-20-one. Neuropharmacology. 43: 1339–1350.

Hosie AM, Clarke L, da Silva HMA, Smart TG. 2009. Conserved site for neurosteroid modulation of GABA_A_ receptors. Neuropharmacology. 56: 149–154.

Hosie AM, Wilkins ME, da Silva HMA, Smart TG. 2006. Endogenous neurosteroids regulate GABA_A_ receptors through two discrete transmembrane sites. Nature. 444: 486–489.

Keller AF, Breton JD, Schlichter R, Poisbeau P. 2004. Production of 5α-reduced neurosteroids is developmentally regulated and shapes GABA_A_ miniature IPSCs in lamina II of the spinal cord. J Neurosci. 24: 907–915.

Khisti RT, Penland SN, VanDoren MJ, Grobin AC, Morrow AL. 2002. GABAergic neurosteroid modulation of ethanol actions. World J Biol Psychiatry. 3: 87–95.

Knabl J, Witschi R, Hosl K, Reinold H, Zeilhofer UB, Ahmadi S, Brockhaus J, Sergejeva M, Hess A, Brune K, Fritschy JM, Rudolph U, Mohler H, Zeilhofer HU. 2008. Reversal of pathological pain through specific spinal GABA_A_ receptor subtypes. Nature. 451: 330–334.

Laurie DJ, Seeburg PH, Wisden W. 1992a. The distribution of 13 GABA_A_ receptor subunit mRNAs in the rat brain. II. Olfactory bulb and cerebellum. J Neurosci. 12: 1063–1076.

Laurie DJ, Wisden W, Seeburg PH. 1992b. The distribution of thirteen GABA_A_ receptor subunit mRNAs in the rat brain. III. Embryonic and postnatal development. J.Neurosci. 12: 4151–4172.

Laverty D, Thomas P, Field M, Andersen OJ, Gold MG, Biggin PC, Gielen M, Smart TG. 2017. Crystal structures of a GABA_A_-receptor chimera reveal new endogenous neurosteroid-binding sites. Nature Structural & Molecular Biology. 24: 977–985.

Lister RG. 1987. The use of a plus-maze to measure anxiety in the mouse. Psychopharmacology (Berl). 92: 180–185.

Low K, Crestani F, Keist R, Benke D, Brunig I, Benson JA, Fritschy JM, Rulicke T, Bluethmann H, Mohler H, Rudolph U. 2000. Molecular and neuronal substrate for the selective attenuation of anxiety. Science. 290: 131–134.

Luisi S, Petraglia F, Benedetto C, Nappi RE, Bernardi F, Fadalti M, Reis FM, Luisi M, Genazzani AR. 2000. Serum allopregnanolone levels in pregnant women: changes during pregnancy, at delivery, and in hypertensive patients. J Clin Endocrinol.Metab. 85: 2429–2433.

Maguire J and Mody I. 2007. Neurosteroid Synthesis-Mediated Regulation of GABA_A_ Receptors: Relevance to the Ovarian Cycle and Stress. Journal of Neuroscience. 27: 2155–2162.

McKernan RM, Rosahl TW, Reynolds DS, Sur C, Wafford KA, Atack JR, Farrar S, Myers J, Cook G, Ferris P, Garrett L, Bristow L, Marshall G, Macaulay A, Brown N, Howell O, Moore KW, Carling RW, Street LJ, Castro JL, Ragan CI, Dawson GR, Whiting PJ. 2000. Sedative but not anxiolytic properties of benzodiazepines are mediated by the GABA_A_ receptor α1 subtype. Nat.Neurosci. 3: 587–592.

Mellon SH and Griffin LD. 2002. Neurosteroids: biochemistry and clinical significance. Trends Endocrinol.Metab. 13: 35–43.

Miller PS, Scott S, Masiulis S, De Colibus L, Pardon E, Steyaert J, Aricescu AR. 2017. Structural basis for GABA_A_ receptor potentiation by neurosteroids. Nature Structural & Molecular Biology. 24: 986–992.

Mohler H. 2012. The GABA system in anxiety and depression and its therapeutic potential. Neuropharmacology. 62: 42–53.

Nusser Z, Sieghart W, Benke D, Fritschy J-M, Somogyi P. 1996. Differential synaptic localization of two major *y*-aminobutyric acid type A receptor α subunits on hippocampal pyramidal cells. Proceedings of the National Academy of Sciences, USA. 93: 11939–11944.

Nusser Z and Mody I. 2002. Selective Modulation of Tonic and Phasic Inhibitions in Dentate Gyrus Granule Cells. Journal of Neurophysiology. 87: 2624–2628.

Olsen RW and Sieghart W. 2009. GABA_A_ receptors: subtypes provide diversity of function and pharmacology. Neuropharmacology. 56: 141–148.

Otis TS and Mody I. 1992. Modulation of decay kinetics and frequency of GABA_A_ receptor-mediated spontaneous inhibitory postsynaptic currents in hippocampal neurons. Neuroscience. 49: 13–32.

Patel R, Bauer CS, Nieto-Rostro M, Margas W, Ferron L, Chaggar K, Crews K, Ramirez JD, Bennett DL, Schwartz A, Dickenson AH, Dolphin AC. 2013. α2δ-1 gene deletion affects somatosensory neuron function and delays mechanical hypersensitivity in response to peripheral nerve damage. J Neurosci. 33: 16412–16426.

Paul SM and Purdy RH. 1992. Neuroactive steroids. FASEB J. 6: 2311–2322.

Pirker S, Schwarzer C, Wieselthaler A, Sieghart W, Sperk G. 2000. GABA_A_ receptors: immunocytochemical distribution of 13 subunits in the adult rat brain. Neuroscience. 101: 815–850.

Poisbeau P, Patte-Mensah C, Keller AF, Barrot M, Breton JD, Luis-Delgado OE, Freund-Mercier MJ, Mensah-Nyagan AG, Schlichter R. 2005. Inflammatory pain upregulates spinal inhibition via endogenous neurosteroid production. J Neurosci. 25: 11768–11776.

Porcu P, Sogliano C, Cinus M, Purdy RH, Biggio G, Concas A. 2003. Nicotine-induced changes in cerebrocortical neuroactive steroids and plasma corticosterone concentrations in the rat. Pharmacol Biochem Behav. 74: 683–690.

Porsolt RD, Le PM, Jalfre M. 1977. Depression: a new animal model sensitive to antidepressant treatments. Nature. 266: 730–732.

Prenosil GA, Schneider Gasser EM, Rudolph U, Keist R, Fritschy JM, Vogt KE. 2006. Specific subtypes of GABA_A_ receptors mediate phasic and tonic forms of inhibition in hippocampal pyramidal neurons. J.Neurophysiol. 96: 846–857.

Puia G, Mienville JM, Matsumoto K, Takahata H, Watanabe H, Costa E, Guidotti A. 2003. On the putative physiological role of allopregnanolone on GABA_A_ receptor function. Neuropharmacology. 44: 49–55.

Purdy RH, Morrow AL, Moore PH, Jr., Paul SM. 1991. Stress-induced elevations of γ-aminobutyric acid type A receptor-active steroids in the rat brain. Proc.Natl.Acad.Sci.U.S.A. 88: 4553–4557.

Reddy DS. 2010. Neurosteroids: Endogenous Role in the Human Brian and Therapeutic Potentials. Progress in brain research. 186: 113–137.

Reeve AJ, Dickenson AH, Kerr NC. 1998. Spinal effects of bicuculline: modulation of an allodynia-like state by an A1-receptor agonist, morphine, and an NMDA-receptor antagonist. Journal of Neurophysiology. 79: 1494–1507.

Rodgers RJ and Johnson NJ. 1998. Behaviorally selective effects of neuroactive steroids on plus-maze anxiety in mice. Pharmacol Biochem Behav. 59: 221–232.

Rudolph U, Crestani F, Benke D, Brunig I, Benson JA, Fritschy JM, Martin JR, Bluethmann H, Mohler H. 1999. Benzodiazepine actions mediated by specific γ-aminobutyric acid_A_ receptor subtypes. Nature. 401: 796–800.

Rupprecht R, Rammes G, Eser D, Baghai TC, Schule C, Nothdurfter C, Troxler T, Gentsch C, Kalkman HO, Chaperon F, Uzunov V, McAllister KH, Bertaina-Anglade V, La Rochelle CD, Tuerck D, Floesser A, Kiese B, Schumacher M, Landgraf R, Holsboer F, Kucher K. 2009. Translocator protein (18 kD) as target for anxiolytics without benzodiazepine-like side effects. Science. 325: 490–493.

Sarkar J, Wakefield S, MacKenzie G, Moss SJ, Maguire J. 2011. Neurosteroidogenesis Is Required for the Physiological Response to Stress: Role of Neurosteroid-Sensitive GABA_A_ Receptors. The Journal of Neuroscience. 31: 18198–18210.

Sigel E and Steinmann ME. 2012. Structure, Function, and Modulation of GABA_A_ Receptors. Journal of Biological Chemistry. 287: 40224–40231.

Smith KS and Rudolph U. 2012. Anxiety and depression: Mouse genetics and pharmacological approaches to the role of GABA_A_ receptor subtypes. Neuropharmacology. 62: 54–62.

Sperk G, Schwarzer C, Tsunashima K, Fuchs K, Sieghart W. 1997. GABA_A_ receptor subunits in the rat hippocampus I: immunocytochemical distribution of 13 subunits. Neuroscience. 80: 987–1000.

Strous RD, Maayan R, Weizman A. 2006. The relevance of neurosteroids to clinical psychiatry: From the laboratory to the bedside. European Neuropsychopharmacology. 16: 155–169.

Tochiki KK, Cunningham J, Hunt SP, Geranton SM. 2012. The expression of spinal methyl-CpG-binding protein 2, DNA methyltransferases and histone deacetylases is modulated in persistent pain states. Mol Pain. 8: 14.

Tretter V, Jacob TC, Mukherjee J, Fritschy JM, Pangalos MN, Moss SJ. 2008. The Clustering of GABA_A_ Receptor Subtypes at Inhibitory Synapses is Facilitated via the Direct Binding of Receptor α2 Subunits to Gephyrin. Journal of Neuroscience. 28: 1356–1365.

Uzunov DP, Cooper TB, Costa E, Guidotti A. 1996. Fluoxetine-elicited changes in brain neurosteroid content measured by negative ion mass fragmentography. Proc.Natl.Acad.Sci U.S.A. 93: 12599–12604.

VanDoren MJ, Matthews DB, Janis GC, Grobin AC, Devaud LL, Morrow AL. 2000. Neuroactive steroid 3alpha-hydroxy-5alpha-pregnan-20-one modulates electrophysiological and behavioral actions of ethanol. J.Neurosci. 20: 1982–1989.

Wieland S, Lan NC, Mirasedeghi S, Gee KW. 1991. Anxiolytic activity of the progesterone metabolite 5 alpha-pregnan-3 alpha-o1-20-one. Brain Res. 565: 263–268.

Wisden W, Laurie DJ, Monyer H, Seeburg PH. 1992. The distribution of 13 GABA_A_ receptor subunit mRNAs in the rat brain. I. Telencephalon, diencephalon, mesencephalon. J.Neurosci. 12: 1040–1062.

Zeilhofer HU, Mohler H, Di Lio A. 2009. GABAergic analgesia: new insights from mutant mice and subtype-selective agonists. Trends in Pharmacological Sciences. 30: 397–402.

Zeilhofer HU, Wildner H, Yevenes GE. 2012. Fast Synaptic Inhibition in Spinal Sensory Processing and Pain Control. Physiological Reviews. 92: 193–235.

